# Visualizing extracellular vesicle-mediated RNA transfer using a novel metabolic labeling approach

**DOI:** 10.1101/2025.08.28.672797

**Authors:** Willemijn S. de Voogt, Tanja Edelbacher, Jerney J.J.M. Francois, Simone Smits, Annet van Wesel, Richard W. Wubbolts, Pieter Vader

## Abstract

Extracellular vesicles (EVs) are key players in intercellular communication, facilitated by the transfer of RNA and other molecular cargo between cells. However, how EV-RNA is processed in recipient cells remains poorly understood, particularly the uptake and intracellular trafficking pathways leading to functional transfer remains elusive. Visualizing EV-RNA in recipient cells may help to address these knowledge gaps. Although EV labeling methods are well-established, tracking endogenous RNA cargo within EVs and in recipient cells is challenging due to its low abundance, requiring highly sensitive labeling methods. Existing RNA labeling methods often lack sensitivity, specificity for donor cell-derived RNA, or are limited to labeling long RNA species. In this study, we propose a novel metabolic labeling approach to track RNA in donor cells, EVs and recipient cells using 5-ethynyl uridine (5-EU). 5-EU is a nucleoside analog of uridine that is incorporated into nascent RNA and can be detected through click chemistry with a fluorescent azide. To enhance RNA labeling efficiency, we overexpressed Uridine-Cytidine Kinase 2 (UCK2) in MDA-MB-231 donor cells, which increased 5-EU incorporation into nascent RNA chains. This approach allowed visualization and quantification of RNA in isolated EVs. Furthermore, co-culturing MDA-MB-231 UCK2^+^ donor cells expressing a CD63-HaloTag fusion protein with HMEC1-eGFP^+^ recipient cells enabled visualization and quantification of intercellular transfer of EVs and their endogenous RNA cargo. This novel approach offers a sensitive and specific tool for studying physiological EV-RNA transfer, thereby advancing our understanding of the biological roles of EVs in intercellular communication.

## Introduction

Extracellular vesicles (EVs) are endogenous, cell-derived nanocarriers capable of functionally delivering RNA cargo to recipient cells ^1–3^. In addition to other mechanisms, such as signaling through EV-cell surface interactions, this intercellular RNA transfer facilitates communication between cells ^4^. For instance, cancer cell-derived EVs have been shown to promote epithelial-to-mesenchymal transition and drive metastasis in secondary tissues by delivering various microRNAs ^5–9^. However, how EVs and their RNA cargo are trafficked intracellularly after uptake, and which pathways mediate release of RNA in the cytosol remains poorly understood ^4^. Simultaneous visualization of EVs and EV-RNA in recipient cells could provide valuable insights into their fate and the role of EV-RNA in intercellular communication. There are well-established methods for labeling EVs, including lipid dyes, fluorescent tagging of EV membrane proteins via genetic engineering, and total protein labeling using NHS esters ^10,11^. In contrast, labeling and detecting endogenous RNA cargo in EV is more challenging, primarily due to the low number of RNA molecules packaged in EVs ^12–14^. Thus, a single-molecule sensitive technique is required to achieve sufficient sensitivity for visualizing specific EV-RNAs in recipient cells. Alternatively, labeling total RNA may be an effective visualization strategy ^13^.

Single-molecule sensitivity can be achieved using fluorescence *in situ* hybridization (FISH)-based approaches, such as single-molecule FISH and molecular beacons, or by incorporating fluorogenic aptamers into target RNA sequences through genetic engineering ^15–18^. Here, multiple FISH probes or aptamers per target RNA molecule are required to obtain this level of sensitivity. Consequently, these methods are restricted to labeling long RNAs, which are even less abundant in EVs than small RNAs ^19^. Additionally, EV-RNAs are likely bound to RNA-binding proteins that are essential for their packaging in EVs. Therefore, the target RNA may be less accessible for FISH probes ^20^. As an alternative to single-molecule techniques, total RNA can be labeled in EV donor cells using RNA-specific dyes, such as SYTO RNASelect ^21,22^. However, recipient cell RNA may compete with EV-RNA for the non-covalent SYTO RNASelect binding. Therefore, this approach cannot be used for selective detection of donor cell-derived RNA in recipient cells. A final approach is to transfect fluorescent RNAs into EV donor cells ^23^. Unfortunately, this process can disturb the endo-lysosomal pathway, and transfection reagent remnants can affect RNA uptake in recipient cells. To overcome these limitations, we propose a novel metabolic labeling approach that allows total RNA labeling in EV donor cells, followed by detection in isolated EVs, and in EV recipient cells.

To metabolically label RNA, we employed 5-EU, which is a cell-permeable nucleoside analog of uridine. After consecutive intracellular phosphorylation steps by Uridine-Cytidine Kinase 1 or 2 (UCK1/UCK2) and cellular monophosphate and diphosphate kinases, it is incorporated as a triphosphate into nascent RNA ^24–26^. 5-EU-labeled RNA can then be visualized using click chemistry, where the ethynyl group is conjugated to an azide-coupled fluorophore. Given the low abundance of RNA molecules in EVs, a high-density labeling method is necessary for visualizing RNA in isolated EVs and recipient cells ^12,27^. Interestingly, overexpression of UCK2 has been shown to significantly enhance 5-EU labeling of nascent RNA, suggesting that UCK1 and UCK2 activity is rate-limiting for 5-EU-based RNA labeling ^28^. In this study, we enhanced 5-EU labeling into nascent RNA following UCK2 overexpression and demonstrate that this metabolic labeling approach enables both the visualization and quantification of endogenously loaded 5-EU-labeled RNA in EVs. Furthermore, we show that this method allows for visualization and quantification of EV-mediated RNA transfer upon contact-free co-culturing of MDA-MB-231 UCK2^+^ donor cells, which also expressed fluorescently tagged CD63, with HMEC1-eGFP^+^ recipient cells. This novel technique provides a sensitive and specific labeling tool, enabling the visualization of the intracellular trafficking of EVs and their RNA cargo, thereby advancing our understanding of their processing, the pathways underlying functional delivery, and their resulting physiological effects.

## Methods

### Cell Culture

MDA-MB-231 were cultured in Dulbecco’s Modified Eagle Medium (DMEM) containing high glucose and L-glutamine (Gibco). HMEC-1 cells were cultured in MCDB-131 medium supplemented with 2 mM L-glutamine (Gibco), 10 ng/mL rhEGF (Peprotech) and 50 nM Hydrocortisone (Sigma) on 0.1% gelatin (Sigma) coated flasks and plates. All cell types were cultured in medium supplemented with 10% Fetal Bovine Serum (FBS) (Capri) and in the presence of 100 u/mL penicillin and 100 µg/mL streptomycin (Gibco) at 37 °C, 5% CO_2_. Cells were grown until 70-80% confluency and passaged two times per week.

### Molecular Cloning

CD63 fused to a GS-flexible linker was PCR-amplified using a Phusion High-Fidelity PCR Kit (New England Biolabs) and cloned into a pHAGE2-EF1α-MCS-IRES-BSD vector using NotI and XbaI restriction enzymes. The HaloTag gene with stop codon was synthesized by Integrated DNA Technologies (IDT) and cloned downstream of CD63-GS using XbaI and BamHI restriction enzymes. FLAG-UCK2 was synthesized by IDT and cloned into a pHAGE2-EF1α-MCS-IRES-BSD vector using NotI and BamHI restriction enzymes. FLAG-UCK2-PURO was created by replacing BSD with PURO using NdeI and SbfI restricition enzymes. For this purpose, PURO, was digested from a pHAGE2-EF1α-MCS-IRES-PURO plasmid with NdeI and SbfI. A Quick Ligase Kit (New England Biolabs) was used for all ligation reactions. Sequencing (Macrogen) of all DNA constructs was performed after each cloning step. The gene fragments and primers that were synthesized by IDT are listed in supplementary table 1.

### Lentiviral Production and Stable Cell Line Generation

Lentiviral particles were produced in HEK293T cells by co-transfecting cells with the envelope plasmid pMD2.G and the packaging plasmid psPAX2 (Addgene plasmids #12559 and #12260, respectively, from Didier Trono) along with a pHAGE2 transfer vector containing the gene of interest. Plasmid DNA was used in a 1:1:2 molar ratio and transfected using Polyethyleneimine (PEI) at a concentration of 2 µg PEI per 1 µg plasmid DNA. 24 h after transfection, the culture medium was refreshed, and viral supernatant was collected after an additional 48 hours. The collected supernatant was centrifuged at 500 x *g* for 5 minutes to remove cells and subsequently filtered through a 0.45 µm syringe filter (Sartorius). Processed lentiviral supernatant was either stored at -80 °C or used immediately for cell transduction. For transduction, target cells were incubated with lentiviral supernatant for 24 hours. Following transduction, the medium was replaced with fresh complete medium containing one of the following selection antibiotics as appropriate: blasticidin (5 µg/mL, InvivoGen), puromycin (2 µg/mL, Gibco). HMEC-1 cells stably expressing eGFP were produced as described elsewhere and cultured in the presence of G418 (500 µg/mL, Capricorn Scientific) ^29^. Cells were maintained under selection for stable expression in the presence of the antibiotic thereafter.

### 5-EU labeling and detection in EV donor cells

EV donor cultured in the presence of 0.5mM 5-ethynyl uridine (5-EU, BroadPharm) in DMEM or 0.05% DMSO for 1 hour -20 hours at 37 °C, 5% CO_2_. Subsequently, cells were washed with PBS and fixed in 2-4% PFA for 20 min-1 hour. Cells were blocked and permeabilized with 5% FCS, 0.1% saponin in PBS for 1 hour at RT. 5-EU was detected using Alexa Fluor 647-azide using the Click-iT Cell Reaction Buffer Kit (Thermo Fisher Scientific) according to manufacturer’s protocol. After two quick washes with 0.1% saponin in PBS, cells were stained by immunofluorescence, or imaged immediately using an EVOS M5000 imaging system (Thermo Fisher Scientific). 5-EU signal was quantified per cell using a custom script in ImageJ version 1.54f ^30^. In brief, a 5-EU signal mask was created using Otsu-based thresholding, and cells were separated using a watershed operation. Mean pixel fluorescence intensity was then measured per cell.

### Immunofluorescence

If applicable, after 5-EU labeling and detection using click chemistry, cells were blocked again with 5% FCS, 0.1% saponin in PBS for 1h at RT, and incubated with 1:200 diluted FLAG antibody (F1804, Sigma) in 1% BSA, 0.1% saponin in PBS overnight at 4 °C. After 3 washes with 0.1% saponin in PBS, cells were incubated with 1:1000 diluted Goat anti-Mouse Alexa Fluor-488 antibody (A28175, Thermo Fisher Scientific). Cells were washed again three times with 0.1% saponin in PBS and the nuclei were stained with 1 µg/mL Hoechst33342 (Thermo Fisher Scientific) prior to confocal imaging.

### EV isolation

MDA-MB-231 wildtype cells and stable MDA-MB-231 UCK2^+^ were grown untill 60–70% confluency. Cells were then incubated in fresh OptiMEM (Gibco) for 16–24 hours containing 0.5mM 5-EU or 0.05% DMSO. Conditioned media were collected and centrifuged at 300 x *g* for 5 minutes and subsequently at 2,000 x *g* for 15 minutes to remove cells and cell debris. The supernatant was filtered using 0.45 µm bottle-top filters (Thermo Fisher Scientific) and concentrated to 15 mL using Tangential Flow Filtration (TFF) with Vivaflow 50R 100 kDa cassettes (Sartorius). The media was further concentrated to a final volume of 1 mL using 100 kDa Amicon Ultra-15 Centrifugal filters (Millipore) by centrifugation at 4,000 x *g* for 30–45 minutes. EVs were isolated from the concentrated media through size exclusion chromatography (SEC) on a Tricorn 10/300 column packed with Sepharose 4 Fast Flow resin (Cytiva), operated on an AKTA Start chromatography system (Cytiva). EV-containing fractions were collected, pooled, and further concentrated to the required volume using 100 kDa Amicon Ultra-4 Centrifugal filters (Millipore).

### Western Blotting

Cells and EVs were lysed in 1x RIPA buffer for 30 minutes at 4 °C. Cell lysates were then centrifuged at 14,000 x *g* to pellet and remove cell debris. Protein concentrations were determined using microBCA Protein Assay Kit (Thermo Fisher Scientific) or Pierce BCA Protein Assay Kit (Thermo Fisher Scientific). Samples were prepared by mixing with SDS sample buffer containing 25mM dithiothreitol (DTT), if necessary, followed by incubation at 95°C for 10 minutes. Equal protein amounts of sample were then loaded onto Bolt 4-12% Bis-Tris polyacrylamide gels (Thermo Fisher Scientific) for electrophoresis. Proteins were transferred onto Immobilon-FL polyvinylidene difluoride (PVDF) membranes (Merck Millipore), and the membranes were blocked with a 1:1 mixture of Intercept Blocking Buffer (LI-COR Biosciences) and Tris-buffered saline (TBS). Primary and secondary antibodies (see Supplementary Table 2) were diluted in a 1:1 mixture of Intercept Blocking Buffer and TBS containing 0.1% Tween20 (TBS-T) for staining. Blots were visualized on an Odyssey M imager (LI-COR Biosciences) using the 700nm and 800nm channels.

### Nanoparticle Tracking Analysis (NTA)

Particle size distributions and concentrations of EVs were determined using a Nanosight S500 analyzer (Malvern Panalytical) equipped with s 405nm laser. Samples were diluted to obtain between 30-100 tracks per frame. Per sample, five 30-second videos were acquired with camera level 6. Videos were then analyzed by applying a detection threshold of 6 using the NTA software version 3.4.

### MemGlow labeling of EVs

Isolated EVs were labeled with 200nM MemGlow560 (Cytoskeleton) for 30 minutes at RT. Next, free dye was removed from labeled EVs using SEC on a XK 16/20 column packed with Sepharose CL4b resin (Cytiva), using an AKTA Start chromatography system (Cytiva). Labeled EV-containing fractions were then collected, pooled and again concentrated using 100kDa Amicon Ultra-4 Centrifugal filters (Millipore).

### Direct Stochastic Optical Reconstruction Microscopy (dSTORM)

CD9 antibody (HI9a, BioLegend) was depleted from sodium azide using a 10kDa MWCO filter ultrafiltration vial (Biotium. 10µg of antibody was conjugated at 4°C overnight with 40 µg/mL NHS-ATTO 488 (Thermo Fisher Scientific) in fresh 100mM borate buffer (pH 8.6). Free NHS-ATTO 488 was subsequently removed using 0.5mL 7kDa MWCO Zeba Dye and Biotin Removal Spin Columns (Thermo Fisher Scientific). CD9-ATTO488 was used immediately or stored at 4°C.

Coverslips (#1.5H, 24 mm x 50 mm, Marienfeld) were cleaned by two rounds of 30 minutes ultrasonication in MilliQ water, followed by 30 minutes ultrasonication in 1M potassium hydroxide. Next, they were dried with nitrogen gas and 50-well CultureWell gaskets (Grace Bio-Labs) were attached. Independent wells on the coverslips were coated with 0.01% poly-L-lysine (PLL) at 37°C for 2 hours. MemGlow 560-labeled EVs were added to the wells overnight at 4°C. The following day, EVs were blocked with 2.5% BSA in PBS for 1 hour at room temperature. 5-EU was subsequently detected using the Click-iT Cell Reaction Buffer Kit (Thermo Fisher Scientific) according to manufacturer’s protocol. Afterwards EVs were blocked in 5% BSA in PBS and stained overnight at 4°C with 10 µg/mL CD9-ATTO488. After staining, the coverslips were washed twice with PBS and imaged the same day in freshly prepared OxEA buffer (pH 8-8.5) ^31^.

dSTORM imaging was performed using a Nanoimager-S (ONI) equipped with 405nm (150 mW), 488nm (1W), 561nm (500 mW), and 640nm (1W) lasers. A 100x objective with a 1.4 numerical aperture (Olympus) was used for imaging. Prior to acquisition, channel alignment was performed with a microscopy slide containing 0.1 µm TetraSpeck™ beads (Thermo Fisher Scientific). Imaging was carried out under TIRF illumination, capturing 2500 frames at 640nm (laser power 30-35%) and 480nm (laser power 30-35%) with a 30ms exposure time. In addition, 500 frames were acquired at 561nm (laser power 30-35%) with a 30ms exposure time to detect the MemGlow 560 signal as a reference.

### Cluster Analysis

Data analysis, including localization filtering and HDBSCAN cluster analysis, was performed using the collaborative discovery (CODI) online platform (https://alto.codi.bio/, ONI). Localization in the independent channels were filtered on frame index. The following filters were applied to all channels: Drift correction: DME; photon count: > 300; localization precision: 0-20 nm. The following for 488nm and 640nm channels: p-value: >1 x 10^-6. The HDBSCAN clustering parameters were set as: area < 80,000 nm^2^; skew < 2.6, and kappa > 23 nm. A 120 nm radius was applied for counting localizations in individual clusters, whereby clusters with > 2 localizations in the 488nm and 640nm channel were considered positive. The percentage of AF647+/MemGlow560+ clusters was calculated as a proportion of the total number of MemGlow 560+ clusters.

### Co-cultures and culture-well insert experiments

MDA-MB-231 UCK2^+^ or UCK2^+^/CD63-Halo^+^ donor cells were seeded as follows: for direct co-culture experiments, 40,000 cells were seeded in a 96-well plate, 120,000 cells in an 8-well Ibidi glass bottom plate, or 240,000 cells in a 24-well plate. Alternatively, silicon 2-well culture-Inserts (Ibidi) were placed in the center of a glass bottom 35µm dish (Ibidi) and 400,000 donor cells were seeded in 400 µL DMEM outside the culture inserts. 16 hours after seeding of donor cells, the medium was replaced with fresh DMEM containing 0.5mM 5-EU (BroadPharm) or 0.05% DMSO. After 8 h of treatment, wells were washed with PBS and HMEC-1 GFP expressing recipient cells were seeded in a 2:1 donor:recipient cell ratio for direct co-cultures. For culture-well insert experiments, inner wells were coated with 0.1% gelatin and 14,000 HMEC-1 GFP recipient cells were seeded. Next, donor cells were washed with PBS and culture well inserts were carefully removed leaving a 500 µm gap in between cell types. Fresh DMEM was added so that donor and recipient cells shared the medium. In addition, DMEM was supplemented with Janelia Fluor 549 HaloTag ligand (Promega) when MDA-MB-231 UCK2^+^/CD63-Halo^+^ donor cells were used. Donor and recipient cells were co-cultured for 16 hours, and, subsequently, trypsinized and fixed in 4% paraformaldehyde (PFA). Cells were then blocked and permeabilized in 5% FBS, 0.3% Triton X-100 in PBS followed by 5-EU detection with Alexa Fluor 647-azide through click chemistry utilizing the Click-iT Cell Reaction Buffer Kit (Thermo Fisher Scientific) according to manufacturer’s protocol. Cells were washed twice with 0.1% Triton X-100 in PBS and were analyzed in 1% FBS in PBS using a FACSCanto II flow cytometer or in PBS by fluorescence microscopy. Flow cytometry analysis was performed using FlowJo v10.8.1.

### Confocal microscopy

Fluorescence imaging of the culture well insert experiments was performed on an Olympus/Evident Spin IXplore SR10 Spinning Disc Confocal Microscope, with 100x UPLXAPO oil-immersion objective (Evident) with Numerical Aperture 1.45. Imaging was performed using the following settings: 640nm (Obis, 100 mW), 50% laser power, 250 milliseconds exposure, BP685/40; 561nm (Obis, 100 mW), 40% laser power, 250 milliseconds exposure, BP 607/36; 488nm (Obis, 100 mW), 20% laser power, 100 milliseconds exposure, BP525/50; 405nm (Obis, 50 mW), 20% laser power, 150 milliseconds exposure, BP447/60. The main dichroid mirror was a quad band (D405/488/561,640). For co-culture experiment, images were analyzed in ImageJ version 1.54f ^30^ using a custom script. In brief, AF647 and Janelia Fluor 549 signals were thresholded to generate binary images. In addition, any signal outside recipient cells (eGFP) was removed. The number of particles in each channel, as well as overlapping objects were quantified.

## Results

### A novel metabolic labeling approach using 5-EU allows visualization of nascent RNA in MDA-MB-231 donor cells and in MDA-MB-231 derived EVs

To label total endogenous RNA in EV donor cells, we used 5-EU, a nucleoside analog of uridine. 5-EU is a cell-permeable compound that differs from endogenous uridine by an ethynyl group at the 5th position of the pyrimidine ring (**Fig. 1A**). After 5-EU treatment of cells, 5-EU is phosphorylated intracellularly after which 5-EU triphosphate can be incorporated by RNA polymerases into nascent RNA chains. We detected 5-EU-labeled RNA using click chemistry with an Alexa Fluor 647 (AF647) – azide (**Fig. 1B**). We first validated 5-EU labeling in MDA-MB-231 EV donor cells. Treating donor cells with 0.5 mM 5-EU enabled specific labeling and detection of nascent RNA, whereas control cells that were treated with 0.05% DMSO showed no signal (**Fig 1C**).

**Figure 1.**
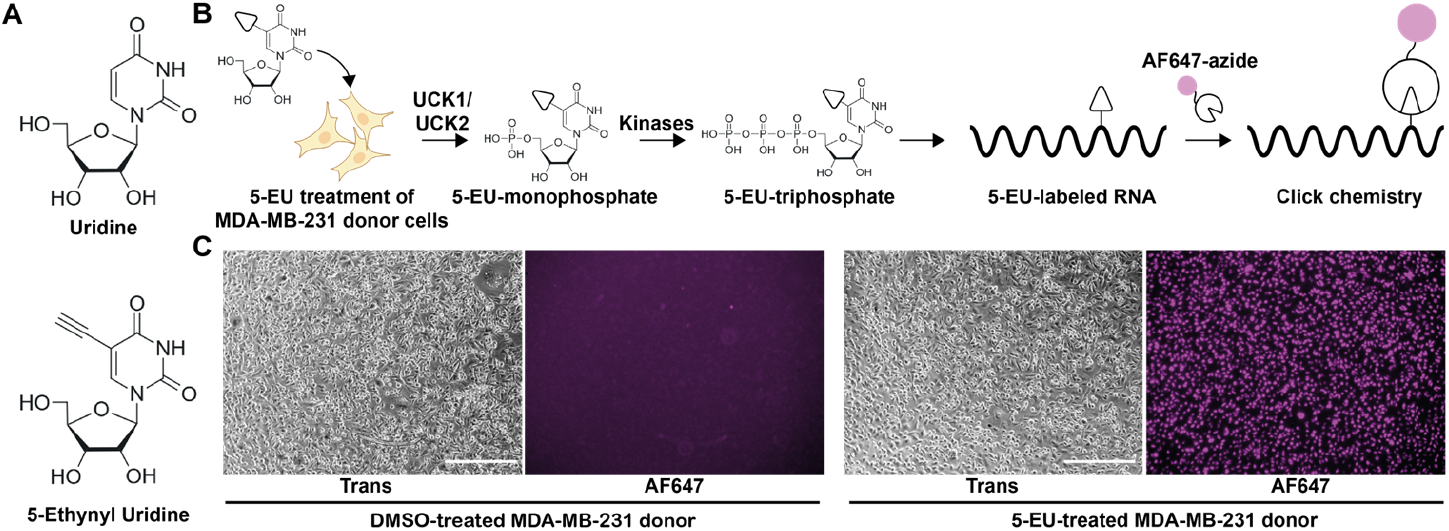
5-EU labels nascent RNA in MDA-MB-231 donor cells and is detected through click chemistry. (A) 5-ethynyl uridine is an RNA-specific analog of endogenous uridine, modified with an ethynyl group at the 5^th^ position of the pyrimidine ring. (B) Upon MDA-MB-231 donor cell treatment, 5-EU is intracellularly phosphorylated by UCK1 or UCK2, and other cellular kinases. The resulting 5-EU triphosphate is then incorporated into newly synthesized RNA. Next, the ethynyl group enables fluorescent detection via a click chemistry reaction with Alexa Fluor 647-azide. (C) Brightfield and fluorescence images demonstrating 5-EU-labeled RNA (magenta) detection in MDA-MB-231 donor cells (right) compared to control cells treated with 0.05% DMSO (left). Scale bars: 300 µm.

Next, to determine whether 5-EU labeled RNA is loaded into EVs and detectable, we isolated EVs from 5-EU-treated MDA-MB-231 donor cells and performed direct stochastic optical reconstruction microscopy (dSTORM) for single EV analysis (**Fig 2**). We successfully isolated EVs using tangential flow filtration followed by size exclusion chromatography, as confirmed by western blot and nanoparticle tracking analysis (NTA) (**Fig 2A, B**). Western blot showed presence of EV marker TSG101, and enrichment of CD9. Additionally, endoplasmic reticulum marker Calnexin was absent in the EV lysate supporting the purity of isolated EVs. NTA revealed a typical EV size distribution profile with a mode size of approximately 115 nm. Using dSTORM and quantitative cluster analysis, we detected CD9 and 5-EU in single EVs and quantified the percentage of 5-EU^+^ EVs. Notably, most EVs were single positive for CD9 (**Fig 2C**). Indeed, cluster analysis revealed that only 2% of EVs stained positive for 5-EU (**Fig 2D**).

**Figure 2.**
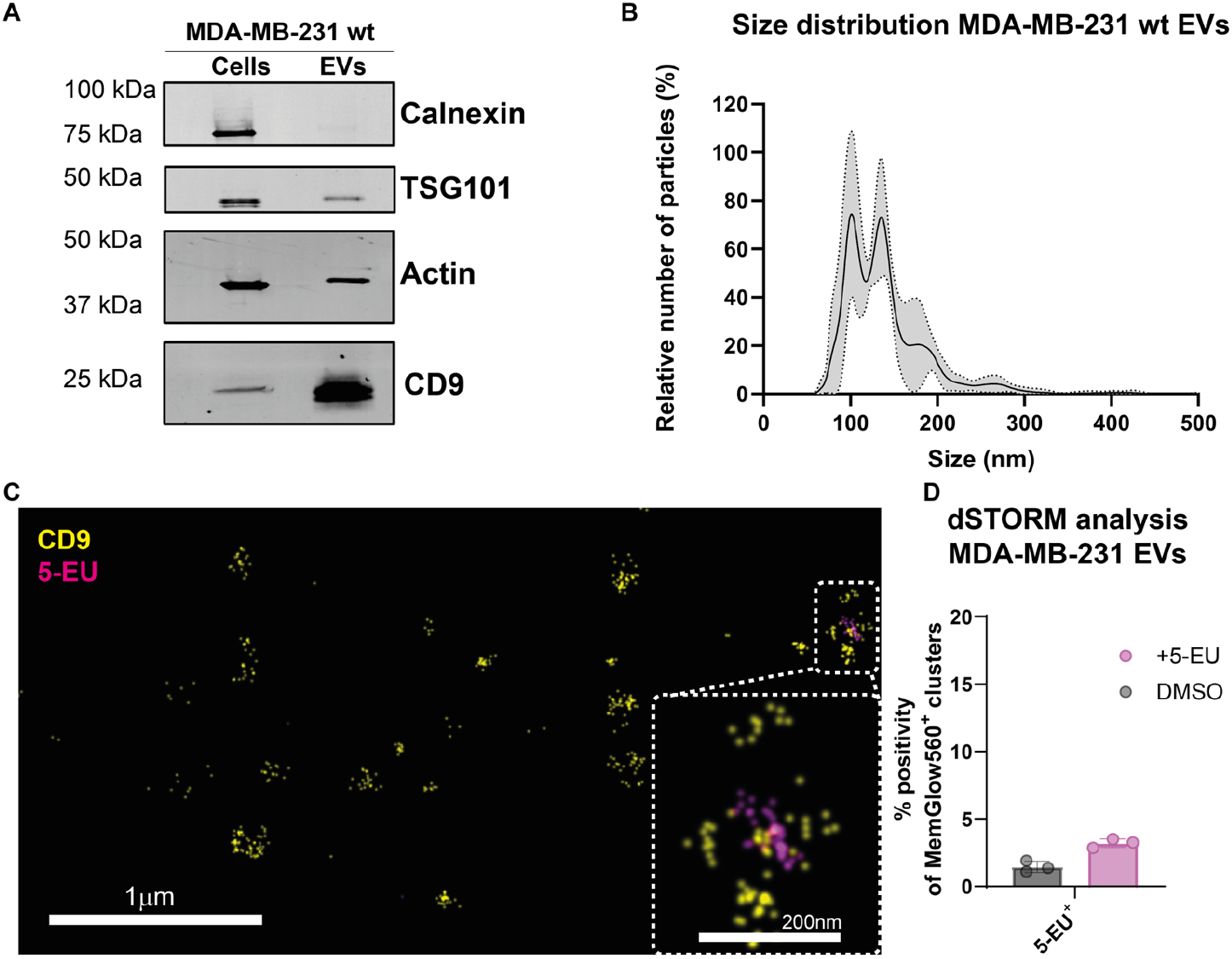
5-EU treatment of donor cells allows for nascent RNA visualization and quantification in donor cell-derived EVs. (A) Western blot analysis of MDA-MB-231 cell and EV lysates comparing common EV marker**s**, TSG101 and CD9, organelle marker Calnexin, and Actin loading control. Equal amounts of protein were loaded for cells and EVs. (B) Size distribution of isolated MDA-MB-231 EVs as determined by NTA. Data is represented as mean ± SD (grey). (C) dSTORM image from a representative experiment showing detection of CD9 (yellow) and 5-EU (magenta) in isolated MDA-MB-231 EVs after incubation of donor cells with 5-EU. Scale bar = 1µm. Bottom right: zoomed-in view of one cluster showing coincidence of CD9 and 5-EU. Scale bar = 200nm. (D) Quantitative cluster analysis of dSTORM data. Results are expressed as the percentage of 5-EU^+^ clusters relative to MemGlow560^+^ clusters and compared to EVs isolated from DMSO-treated donor cells.

### UCK2 overexpression improves labeling density of 5-EU in RNA in EV donor cells and isolated EVs

It is likely that the apparent relatively low percentage of 5-EU-positive EVs detected (**Fig 2D**) is an underestimation of the true percentage of RNA-positive EVs, as a result of low RNA 5-EU labeling efficiency. We hypothesized that this may have been caused by the fact that 5-EU phosphorylation by UCK1 or UCK2 may be outcompeted by endogenous uridine. To improve 5-EU phosphorylation in EV donor cells and enhance RNA labeling, we generated a MDA-MB-231 donor cell line stably overexpressing UCK2 (**Fig 3A, B**). We validated the expression of FLAG-UCK2 by western blot and immunofluorescence (**Fig 3C, D**). Importantly, when we compared RNA labeling using fluorescence microscopy in MDA-MB-231 wildtype donor cells and MDA-MB-231 UCK2^+^ donor cells after 1, 2, 4, and 20 hours of 5-EU labeling, we observed a clear increase in 5-EU signal indicating more efficient RNA labeling in UCK2^+^ donor cells (**Fig 3E**). Quantification of the fluorescent signal per cell over time showed that nascent RNA labeling in wildtype and UCK2^+^ cells demonstrated a similar kinetic profile. However, for MDA-MB-231 UCK2^+^ donor cells, a higher maximum fluorescence signal was obtained (**Fig. 3F**). This suggests that more efficient labeling of 5-EU in these cells is a result of a higher labeling density of 5-EU in RNA. To determine if UCK2 overexpression in donor cells increased the observed proportion of 5-EU^+^ EVs, we isolated EVs from MDA-MB-231 UCK2^+^ donor cells and performed super-resolution microscopy by dSTORM. Indeed, we detected 5-EU in approximately 8% of the MDA-MB-231 UCK2^+^ EVs, which is remarkably higher than the percentage of 5-EU^+^ EVs derived from MDA-MB-231 wildtype cells (**Fig 3G, H**).

**Figure 3.**
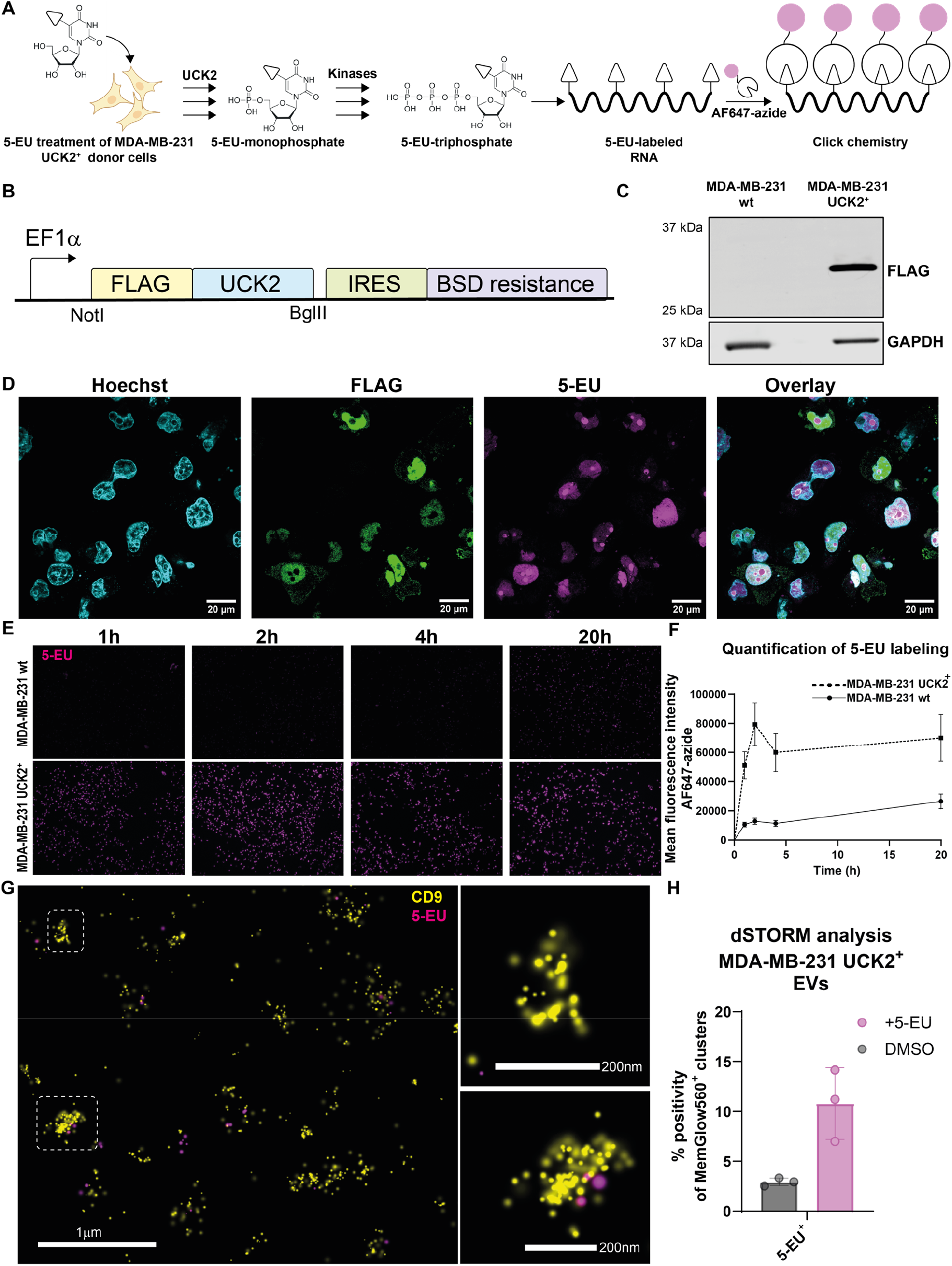
Overexpression of UCK2 in MDA-MB-231 donor cells enhances 5-EU labeling of RNA and detection in cells and isolated EVs. (A) Overexpression of UCK2 in donor cells enhances the 5-EU to 5-EU monophosphate conversion. As a result, more 5-EU monophosphate is converted to 5-EU triphosphate by UMP-CMPK and NDPK, and a higher ethynyl labeling density and more sensitive detection is obtained using click chemistry. Created in BioRender. Vader, P. (2025) https://BioRender.com/3pwgjvv Lentiviral DNA construct engineered for stable overexpression of UCK2 in donor cells. UCK2 is tagged with a FLAG-tag enabling detection of UCK2 expression. Additionally, an internal ribosomal entry site (IRES) is encoded upstream of a blasticidin (BSD) resistance gene for generating and maintaining stable expression of UCK2. (C) Western blot analysis for FLAG and GAPDH expression in MDA-MB-231 UCK2^+^ cell lysate and MDA-MB-231 wildtype cell lysate. Equal amounts of protein were loaded. (D) Confocal microscopy image of MDA-MB-231 UCK2^+^ donor cells. FLAG expression (green) was detected through immunofluorescence, 5-EU (magenta) through click chemistry, and nuclei (cyan) using Hoechst33342. (E) Fluorescence microscopy images showing a comparison of 5-EU labeling (magenta) in MDA-MB-231 UCK2^+^ and MDA-MB-231 wildtype donor cells upon incubation with 5-EU for 1 hour, 2 hours, 4 hours, and 20 hours. (F) Quantification of fluorescent signal from microscopy images in E. Mean fluorescence intensity per cell is represented over time for wildtype donor cells and UCK2+ donor cells. Data is presented as mean ± SD (G) Representative dSTORM image of EVs isolated from MDA-MB-231 UCK2^+^ after 5-EU treatment. EVs were labeled using CD9 antibodies (yellow) and RNA was labeled using 5-EU (magenta). (H) Quantitative cluster analysis of dSTORM experiment. Data are represented as percentage of 5-EU^+^ clusters/MemGlow560^+^ clusters and compared to EVs isolated from DMSO treated UCK2^+^ donor cells.

### 5-EU labeling in UCK2^+^/CD63-Halo^+^ sMDA-MB-231 donor cells allows visualization and quantification of EV-RNA transfer to recipient cells

To assess whether our metabolic labeling approach allows detection of EV-mediated RNA transfer, we engineered a new MDA-MB-231 donor cell line, stably expressing both UCK2 and CD63-HaloTag (**Fig 4A**). HaloTag (Halo) is an enzyme that covalently binds ligands containing a chloroalkane linker, which can be attached to interchangeable molecules, including bright and cell-permeable fluorophores ^32^. This method enables high-sensitivity labeling of CD63^+^ EVs. We validated the expression of CD63-Halo and UCK2 via western blot and confocal microscopy (**Fig 4B, C**). Initially, we performed a direct co-culture experiment in which MDA-MB-231 UCK2^+^/CD63-Halo^+^ donor cells and HMEC-1 GFP^+^ recipient cells were randomly distributed and cultured together in a single well. Strikingly, we observed a high and UCK2-dependent uptake of 5-EU in recipient cells using flow cytometry (**Fig S1A-C**). However, microscopy analysis revealed that most of the 5-EU signal in recipient cells localized to the nucleus, suggesting a non-EV-mediated pathway of 5-EU transfer (**Fig S1D**).

**Figure 4.**
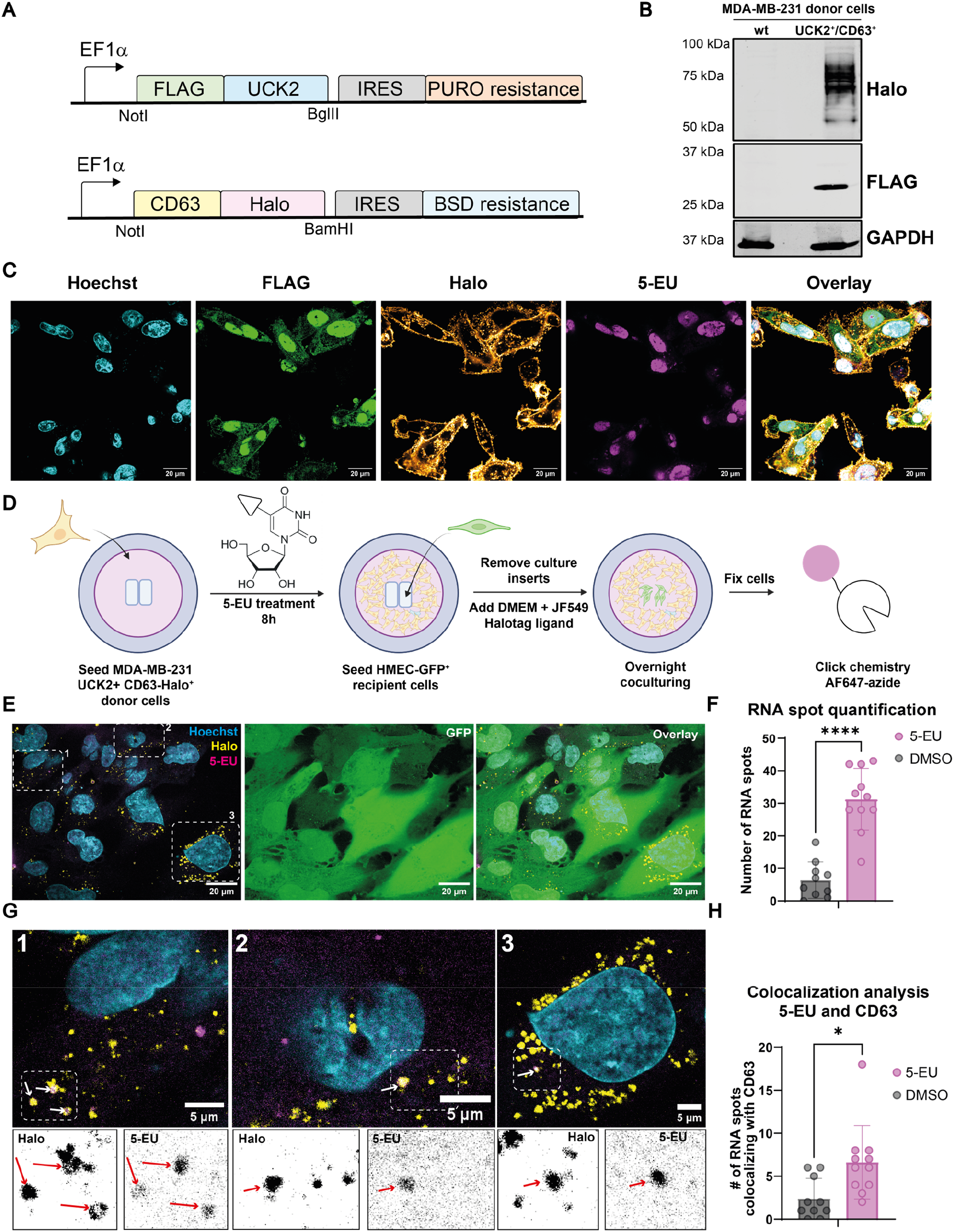
5-EU labeling in UCK2+/CD63-Halo+ MDA-MB-231 donor cells allows detection and quantification of EV-mediated RNA transfer. (A) Lentiviral DNA constructs engineered for stable co-overexpression of UCK2 and CD63-Halo in donor cells. CD63-Halo is expressed as a fusion construct. Additionally, an IRES UCK2 is tagged with a FLAG-tag enabling detection of UCK2 expression. Additionally, an IRES and BSD resistance gene enable antibiotic selection pressure for generating and maintaining stable cells. A puromycin (PURO) resistance gene has replaced (BSD) in the FLAG-UCK2 expressing constructs. (B) Western blot analysis for FLAG, Halo and GAPDH expression in MDA-MB-231 UCK2^+^ cell lysate and MDA-MB-231 wildtype cell lysate. Equal amounts of protein were loaded. (C) Confocal microscopy image of MDA-MB-231 UCK2^+^ donor cells. FLAG expression (green) was detected through immunofluorescence, Halo expression through Janelia Fluor 549-Halotag ligand (yellow), 5-EU (magenta) through click chemistry, and nuclei (cyan) using Hoechst33342. (D) Experimental set-up of a co-culture with culture well inserts. Silicon culture well inserts were placed in the center of a glass bottom culture dish. MDA-MB-231 UCK2^+^/CD63-Halo^+^ donor cells were cultured in the presence of 5-EU for 8 hours outside the inserts. HMEC-GFP^+^ recipient cells were then seeded within the culture well inserts. Silicon inserts were removed and donor and recipient cells were cultured in shared medium containing Janelia Fluor 549-Halotag ligand. After co-culturing overnight, cells were fixed, and 5-EU was detected using Alexa Fluor 647-azide click chemistry. Created in BioRender. Vader, P. (2025) https://BioRender.com/7i9qete (E) Representative confocal microscopy image of the center of the glass bottom culture dish. Left: Overlay image of nuclei detected using Hoechst33342 (cyan), CD63-Halo (yellow) using Janelia Fluor 549-Halotag ligand, and 5-EU (magenta) using Alexa Fluor 647-azide. Middle: GFP (green) signal as expressed by HMEC-1 recipient cells. Right: Overlay image of all channels. Scale bar = 20µm (F) Quantification of number of RNA spots detected in recipient cells. Co-cultures whereby donor cells were treated with 5-EU were compared to control co-cultures whereby donor cells treated with 0.05% DMSO. Each datapoint represents a single image. Results are shown as means ± SD. Statistical significance was assessed using an unpaired T-test. Indicated p-value: **** = p < 0.0001 (G) Top: Zoom-ins of single cells, outlined in boxed regions in panel E. Numbers correspond to the numbers of each box in e. Coincidence events of CD63-Halo and 5-EU are indicated with white arrows. Scale bar = 5µm. Bottom: Split channel views of boxed regions from top panel displaying Halo and 5-EU signal separately. Coincidence events are indicated with red arrows. (H) Quantification of number of RNA spots per that coincides with CD63-Halo. Co-cultures whereby donor cells were treated with 5-EU were compared to control co-cultures whereby donor cells treated with 0.05% DMSO. Each datapoint represents a single image. Results are shown as means ± SD. Statistical significance was assessed using an unpaired T-test. Indicated p-value: * = p < 0.05.

We hypothesized that in a direct co-culture setup, contact-dependent uptake of phosphorylated 5-EU might occur, possibly via gap junctions, tunneling nanotube formation, or trogocytosis ^33–35^. Therefore, we designed a co-culture assay in which donor and recipient cells are physically separated by silicon culture well inserts. After incubating donor cells with 5-EU, recipient cells were seeded into the inserts. Next, the inserts were removed, creating a 500 µm gap between donor and recipient cells, followed by overnight culturing in shared medium and 5-EU detection via click chemistry (**Fig 4D, S2**). Notably, in this contact-free co-culture set-up, we observed negligible 5-EU signal in the nucleus of recipient cells. At the same time, we still observed clear uptake of CD63-Halo EVs in recipient cells. Furthermore, in individual cells, we observed distinct 5-EU spots, consistent with an endosomal localization pattern as expected upon uptake of RNA-containing EVs. In addition, we were able to quantify this RNA transfer (**Fig 4E, F**). In some instances these 5-EU-RNA spots colocalized with CD63^+^ EVs, suggesting indeed successful detection of EV-mediated RNA transfer (**Fig 4G, H**). Non-coincided spots may represent uptake of CD63^-^ EV-RNA, RNA that has been released from EVs, or RNA that is transferred intercellularly in an EV-independent manner. Together, these data show that RNA labeling in 5-EU overexpressing donor cells allows for visualization and quantification of EV-mediated RNA transfer in recipient cells.

## Discussion

Over the past decades, numerous studies have demonstrated that extracellular vesicles (EVs) can transport and deliver RNA and other molecular cargo to recipient cells, acting as natural mediators of intercellular communication ^1–3,36^. However, the extent to which RNA transfer specifically contributes to EV function, compared to alternative mechanisms such as signaling induced by EV-cell surface interactions, remains unclear ^37^. For example, while many studies report physiological roles for functionally transferred miRNAs, they often fail to demonstrate the EV dependency of this transfer ^38–41^. This discussion is further fueled by the relatively low abundance of RNAs in EVs ^4,12,14,42,43^. Several studies have employed a Cre recombinase reporter system to evaluate EV-mediated transfer of Cre mRNA. However, the possibility of reporter cell activation due to the transfer of functional Cre protein cannot be excluded in this assay ^44,45^. To address this limitation, in previous work we assessed the efficiency of EV-mediated functional single guide RNA (sgRNA) delivery using a CRISPR/Cas9-based reporter system. In co-culture experiments, donor cells expressing sgRNA were paired with recipient reporter cells expressing Cas9 and the target sequence. Reporter cell activation was observed in only 0.2% of recipient cells, indicating limited EV-mediated functional sgRNA delivery, which could be explained by the low number of sgRNA molecules packaged in EVs. Interestingly, reporter cell activation, and thus functional EV-mediated sgRNA delivery, depended on the specific combination of donor and recipient cells ^36^. It is therefore tempting to speculate that RNA delivery and subsequent biological effects are selective and regulated processes rather than passive and stochastic events. A deeper understanding of how EVs and their RNA cargo are processed in recipient cells is essential for further elucidating their biological roles.

To achieve this, advanced techniques that allow simultaneous labeling and visualization of both EVs and their RNA cargo are crucial. To visualize RNA cargo within EVs and in recipient cells, a sensitive, donor cell-RNA specific technique is required. Current RNA labeling methods suffer from limited sensitivity and specificity or require RNA transfection into donor cells or RNA engineering, making them unsuitable for studying endogenous intercellular RNA transfer ^13,15,18^. To achieve RNA visualization, we here developed a novel metabolic labeling approach for total endogenous RNA using 5-ethynyl uridine (5-EU), a nucleoside analog that is incorporated into nascent RNA by cellular machinery ^26^. Upon fluorescent detection via click chemistry, we are able to visualize 5-EU in EV donor cells with high specificity. Interestingly, we observed that the majority of 5-EU signal localized in the nucleus. Indeed, transcription occurs at transcription sites in nuclear domains ^46^. Additionally, we and others observed most intense staining in nucleoli, which are transcription sites of abundant ribosomal RNAs ^26^.

To assess the feasibility of using this metabolic labeling approach to track EV-mediated intercellular RNA transfer, we first aimed to detect 5-EU-labeled RNA in isolated EVs. Loading efficiency of 5-EU-labeled RNA into EVs is likely time-dependent and influenced by several factors. First, nascent RNA must translocate from the nucleus to the cytoplasm and maintain intact before incorporation into EVs. Second, RNA must be packaged into EVs for which timing is affected by kinetics of EV biogenesis and release. Third, there is a dynamic balance between EV secretion and uptake, meaning that pulsed RNA labeling may result in the transient presence of 5-EU-labeled RNA in EVs in the extracellular space. Our limited understanding of the cumulative kinetics of these processes makes it highly challenging to pinpoint optimal timing for 5-EU treatment. In our assays, we used 8-16-hour incubations with 5-EU, however, a better understanding of the kinetics of RNA labeling, release and consumption may enhance 5-EU-labeled RNA levels in EVs.

We used dSTORM imaging to detect RNA in EVs. Initially, only a small subset of EVs (∼2%) stained positive for 5-EU. We hypothesized that the preferential incorporation of uridine over 5-EU limits the number of labels incorporated per RNA molecule, thereby reducing detection efficiency. Previous studies on 18S ribosomal RNA have shown that, on average, one 5-EU is incorporated for every 35 uridines ^26^. Assuming a random distribution of adenosine, cytidine, guanosine, and uridine within RNA sequences, this corresponds to approximately one 5-EU label per 140 nucleotides ^47^. Combined with the low abundance of RNA in EVs, this level of labeling is likely insufficient for effective RNA detection in recipient cells.

In an attempt to enhance labeling density, we overexpressed uridine-cytidine kinase 2 (UCK2) in MDA-MB-231 donor cells. Both UCK1 and UCK2 catalyze the phosphorylation of uridine to its monophosphate form, which is further converted to uridine di- and tri-phosphate by other cellular kinases ^47^. UCK2 overexpression has previously been shown to significantly enhance 5-EU labeling of RNA ^28^. Notably, UCK2 exhibits 15-20 fold higher catalytic activity than UCK1, aligning with its tissue-specific expression patterns ^48^. While UCK1 is broadly expressed in various tissues, UCK2 is primarily found in the human placenta and various tumor cells ^49^. Interestingly, upregulation of UCK2 has been associated with enhanced tumor progression and poor clinical outcomes in, for instance, breast cancer and hepatocellular carcinoma ^50,51^. The potential influence of UCK2 overexpression on the metabolism of endogenous pyrimidines and its broader impact on secreted EV composition and functionality remains to be fully explored. However, in our experiments, UCK2-overexpressing cells did not display any observable signs of toxicity, suggesting that its overexpression does not inherently compromise cell viability under our conditions.

Following 5-EU treatment of MDA-MB-231 UCK2+ donor cells, we observed increased labeling in donor cells and detected 5-EU in approximately 8% of isolated EVs. However, it remains unclear whether this percentage accurately reflects stoichiometry or if it is influenced by labeling and detection efficiencies. Given the heterogeneity of the EV population, which varies in size and subcellular origin, it is possible that RNA may be selectively loaded into specific subsets of EVs ^52^. For example, various RNA motifs have been identified, which have been shown be selectively loaded into exosomes, a subtype of EVs ^18,53,54^. In addition, with increasing EV size, the proportion of EV lumen over membrane increases drastically ^55^. Therefore, it may be that larger EVs contain the vast majority of RNA cargo. Finally, our detection system may favor longer RNAs with a higher number of 5-EU labels, or those with uridine-rich sequences.

Next, we proceeded to evaluate if our approach allows detection of RNA in recipient cells. To achieve this, we utilized a co-culture system to visualize EV-mediated RNA transfer without the need for EV isolation procedures, thereby closely mimicking physiological RNA transfer. Interestingly, in direct co-culture experiments, the majority of transferred 5-EU localized to the nucleus of HMEC-1 recipient cells. This suggests that phosphorylated 5-EU may be transferred via mechanisms beyond EVs and subsequently incorporated into nascent RNA in recipient cells. Phosphorylated forms of 5-EU may be actively secreted by donor cells, as uridine diphosphate and triphosphate are known to be released into the extracellular space through various mechanisms, where they play a key role in signaling via P2Y_6_ receptor activation ^56,57^. However, whether 5-EU di- and triphosphate can in fact be passively or actively taken up by recipient cells is not well understood. Alternative mechanisms, such as transfer via gap junctions, tunneling nanotubes, or trogocytosis may be responsible for mediating the direct transfer of phosphorylated forms of 5-EU or 5-EU-labeled RNA ^33–35^. These findings highlight the complexity of studying EV-mediated RNA transfer.

To rule out cellular contact-dependent 5-EU or RNA transfer in direct co-culture, we performed co-cultures using culture well inserts, ensuring physical separation of donor and recipient cells while allowing EV-mediated intercellular RNA transfer. Although some studies have shown coincidence of EVs and RNA at the single EV level ^15,18^, visualization of EVs and EV-RNA in recipient cells remains uncommon. One study employed a transwell contact-free co-culture system in which PalmGFP^+^ donor cells were transfected with labeled siRNA, revealing low levels of EV and siRNA coincidence ^23^. Donor-cell transfection, however, may alter the endolysosomal system and influence EV release, making this approach less suitable for studying physiological EV-mediated RNA transfer. To enable this, we developed an approach to detect EVs and total, endogenously loaded EV-RNA. We generated an MDA-MB-231 donor cell line overexpressing both UCK2 and CD63-Halo, enabling simultaneous detection of CD63+ EVs and RNA in recipient cells to confirm EV dependency. Our results demonstrate significant intercellular RNA transfer and show that RNA partially coincides with CD63-Halo in recipient cells, supporting an EV-mediated mechanism. RNA that did not coincide with CD63-Halo may have been released from EVs or originate from a distinct EV subtype lacking CD63-Halo. The low number of double CD63^+^ and 5-EU^+^ spots observed aligns with the sparsity of labeling, and therefore with our detection limit. However, acquiring z-stacks instead of single confocal slices, and thus imaging the entire cell volume, may enhance detection. Altogether, we have demonstrated that our approach allows for detection and quantification of EV-mediated endogenous RNA transfer, enabling study of physiological roles of RNA delivery by EVs.

In future work, our novel metabolic labeling approach can be used to investigate the processing of EVs and their RNA content in recipient cells. This method enables the study of RNA and EV coincidence over time, potentially providing insights into the kinetics of RNA delivery. Additionally, coincidence of RNA with key intracellular trafficking components, such as early and late endosomal markers like Rab5 and LAMP1, can be determined as has been previously done for EVs but not for their RNA cargo ^58,59^. This could provide valuable insights into the selective pathways of RNA trafficking prior to release into the recipient cell cytoplasm. Finally, donor cell lines expressing fusion proteins of various EV membrane markers and Halo can be used in combination with UCK2 overexpression and 5-EU labeling. This allows to determine whether different RNAs in various EV subtypes exhibit distinct trafficking patterns and whether these differences are linked to functional outcomes in recipient cells. Unfortunately, our method is not compatible with live-cell imaging, as it requires copper as a catalyst for the click reaction. However, novel nucleoside and nucleotide analogs are being developed, including intrinsically fluorescent cytidine analogs and azide-modified nucleosides ^25,60,61^. The latter can be detected using copper-free click chemistry, potentially enabling live-cell tracking of RNA cargo ^61^. This advancement would provide additional spatiotemporal data, significantly enhancing our understanding of RNA processing dynamics. For optimal labeling efficiency comparable to 5-EU, however, an alternative nucleoside analog must also be a substrate of UCK2. Furthermore, for inherently fluorescent alternatives, it is crucial that they match or exceed the brightness of common dyes, such as Alexa Fluor 647, to maintain the sensitivity of the assay.

To conclude, we developed a novel metabolic labeling approach to track EV-mediated RNA transfer, providing an opportunity to investigate processing of EVs and their RNA cargo in recipient cells. This may provide important insights into the pathways of uptake and intracellular trafficking that lead to RNA delivery and increase our understanding of the physiological role of EV-mediated RNA delivery.

