## Supplementary information for "Visualizing extracellular vesicle-mediated RNA transfer using a novel metabolic labeling approach"

**Supplementary Table 1.** Gene fragment and primer sequences used in this study for generating lentiviral DNA constructs.

| gBlock/primer | Sequence |
|---|---|
| NotI-HaloTag-NsiI | agctgaGCGGCCGCcatggcagaaatcggtactggctttccattcgacccccattatgtggaagtcctgggcgagcgcatgc<br>actacgtcgatgttggtccgcgcgatggcacccctgtgctgttcctgcacggtaacccgacctcctcctacgtgtggcgcaacatcatc<br>ccgcatgttgcaccgacccatcgctgcattgctccagacctgatcggtatgggcaaatccgacaaaccagacctgggttatttcttcg<br>acgaccacgtccgcttcatggatgccttcatcgaagccctgggtctggaagaggtcgtcctggtcattcacgactggggctccgctct<br>gggtttccactgggccaagcgcaatccagagcgcgtcaaaggtattgcatttatggagttcatccgccctatcccgacctgggacga<br>atggccagaatttgcccgcgagaccttccaggccttccgcaccaccgacgtcggccgcaagctgatcatcgatcagaacgtttttatc<br>gagggtacgctgccgatgggtgtcgtccgcccgctgactgaagtcgagatggaccattaccgcgagccgttcctgaatcctgttgac<br>cgcgagccactgtggcgcttcccaaacgagctgccaatcgccggtgagccagcgaacatcgtcgcgctggtcgaagaatacatgg<br>actggctgcaccagtcccctgtcccgaagctgctgttctggggcaccccaggcgttctgatcccaccggccgaagccgctcgcctgg<br>ccaaaagcctgcctaactgcaaggctgtggacatcggcccgggtctgaatctgctgcaagaagacaacccggacctgatcggcag<br>cgagatcgcgcgctggctgtcgacgctcgagatttccggctagATGCATatgcga |
| NotI-FLAG-UCK2-BglII | acggtcGCGGCCGCatggattacaaagacgatgacgataaggccggggacagcgagcagaccctgcagaaccaccagca<br>gcccaacggcggcgagcccttccttataggcgtcagcgggggaacagctagcggcaagtcttccgtgtgtgctaagatcgtgcagct<br>cctggggcagaatgaggtggactatcgccagaagcaggtggtcatcctgagccaggatagcttctaccgtgtccttacctcggagca<br>gaaggccaaagccctgaagggccagttcaactttgaccacccggatgcctttgacaatgaactcattctcaaaacactcaaagaaa<br>tcactgaagggaaaacagtccagatccccgtgtatgactttgtctcccattcccggaaggaggagacagttactgtctatcccgcag<br>acgtggtgctctttgaagggatcctggccttctactcccaggaggtacgagacctgttccagatgaagctttttgtggatacagatgcg<br>gacacccggctctcacgcagagtattaagggacatcagcgagagaggcagggatcttgagcagattttatctcagtacattacgttc<br>gtcaagcctgcctttgaggaattctgcttgccaacaaagaagtatgctgatgtgatcatccctagaggtgcagataatctggtggcca<br>tcaacctcatcgtgcagcacatccaggacatcctgaatggagggccctccaaacggcagaccaatggctgtctcaacggctacacc<br>ccttcacgcaagaggcaggcatcggagtccagcagcaggccgcattgaAGATCTgactgc |
| FW_NotI-CD63 | atcttataGCGGCCGCatggcggtggaaggaggaatgaaatgtg |
| RV_XbaI-GSlinker-CD63 | atgatgtaTCTAGAcGAGCCTCCACCGCCcatcacctcgtagccacttctgatac |

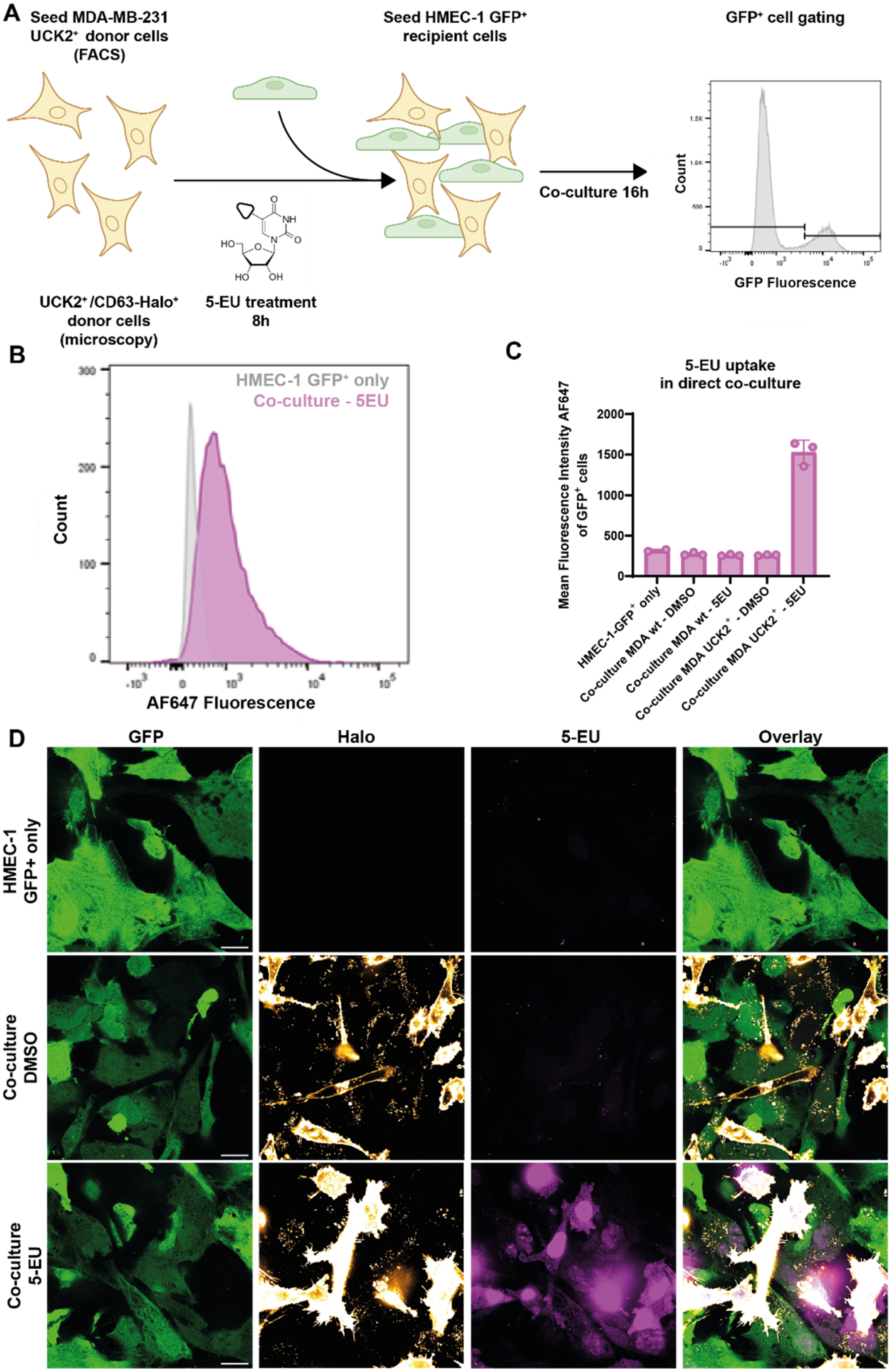

A
Seed MDA-MB-231 UCK2+ donor cells (FACS)
Seed HMEC-1 GFP+ recipient cells
GFP+ cell gating
Co-culture 16h
UCK2+/CD63-Halo+ donor cells (microscopy)
5-EU treatment 8h
Count
GFP Fluorescence
B
HMEC-1 GFP+ only
Co-culture - 5EU
Count
AF647 Fluorescence
C
5-EU uptake in direct co-culture
Mean Fluorescence Intensity AF647 of GFP+ cells
HMEC-1-GFP+ only
Co-culture MDA wt - DMSO
Co-culture MDA wt - 5EU
Co-culture MDA UCK2+ - DMSO
Co-culture MDA UCK2+ - 5EU
D
GFP
Halo
5-EU
Overlay
HMEC-1 GFP+ only
Co-culture DMSO
Co-culture 5-EU

**Supplementary figure S1. In a direct co-culture assay, 5-EU signal in recipient cells localizes to the nucleus.**
(A) MDA-MB-231 donor cells stably expressing UCK2$^+$ (for FACS) or UCK2$^+$/CD63-Halo$^+$ (for confocal microscopy) were treated with 5-EU for 8 hours, washed, and subsequently co-cultured with HMEC-1 GFP$^+$ recipient cells. For FACS analysis, GFP$^+$ recipient cells were gated. Created in BioRender. Vader, P. (2025) https://BioRender.com/yqpwor8 (B) AF647 fluorescence was measured in the GFP$^+$ recipient cell population. The histogram shows AF647 fluorescence in co-cultures with 5-EU-treated UCK2$^+$ donor cells (magenta) compared to the recipient cell-only control (grey). (C) Mean fluorescence intensity (MFI) of AF647 in GFP$^+$ recipient cells under different conditions. Experimental groups included recipient cells alone, co-cultures with MDA-MB-231 wildtype donor cells treated with either DMSO or 5-EU, and co-cultures with MDA-MB-231 UCK2$^+$ donor cells treated with either DMSO or 5-EU. Representative biological replicate is shown. Means ± SD of three individual technical replicates are displayed. (D) Confocal microscopy images showing 5-EU transfer upon co-culturing with MDA-MB-231 UCK2$^+$ donor cells. Upper: recipient cell-only control. Middle: co-culture with DMSO-treated donor cells. Lower: co-culture with 5-EU-treated donor cells. GFP (green), Halo (yellow), and 5-EU (magenta) signals are displayed, along with overlay images. Scale bars: 20 µm.

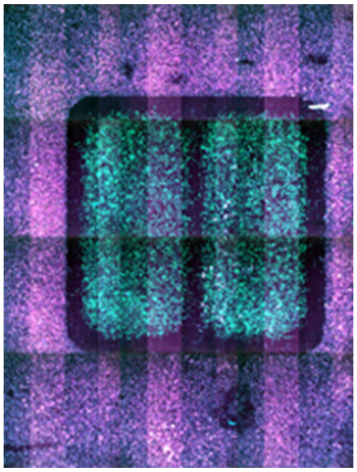

**Supplementary figure S2. Culture setup in which EV donor and recipient cells are physically separated by silicon culture well inserts allows for direct, contact-free co-culture assays.**
Overview image of co-culture 5-EU-treated MDA-MB-231 UCK2$^+$ donor cells (outer) and HMEC-1-GFP$^+$ recipient cells (inner). 5-EU (magenta) and GFP (green) fluorescence are displayed. Donor and recipient cells were separated by 500 µm thick culture well inserts.
